## Supplementary figures and images for "Peptide-based GIP receptor inhibition exhibits modest metabolic changes in mice when administered either alone or combined with GLP-1 agonism"

### supplemental figures

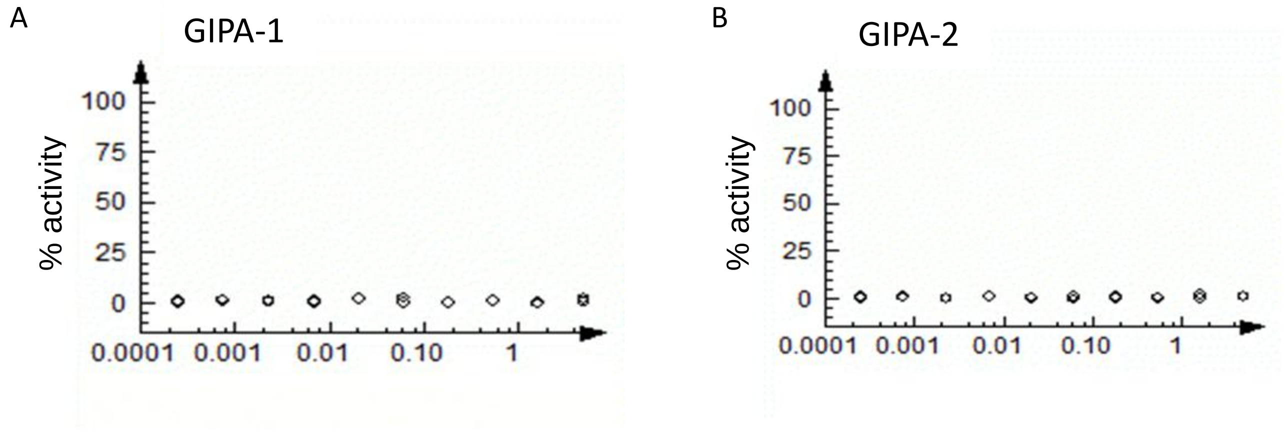

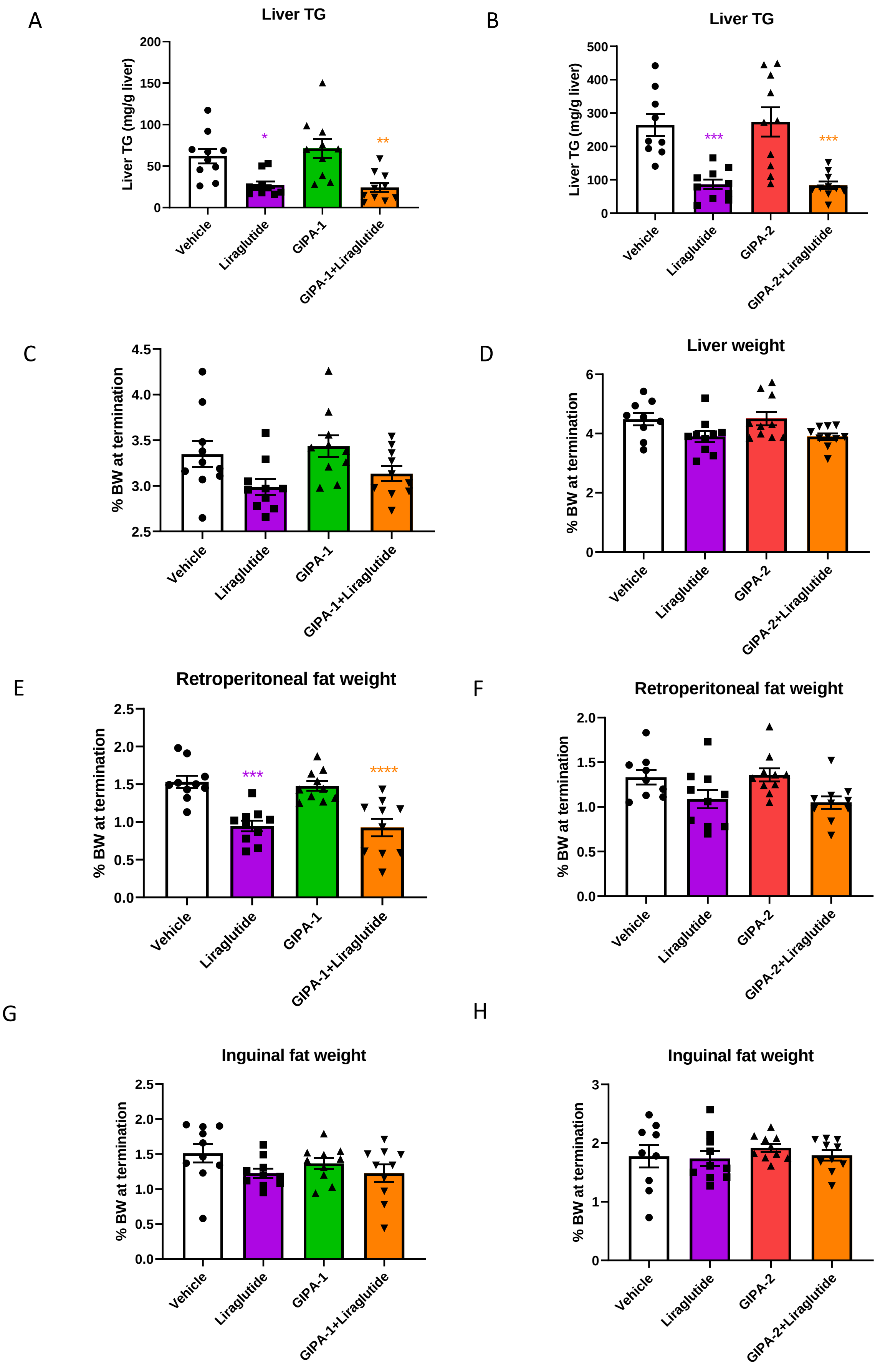
